# SPIRAL: Significant Process InfeRence ALgorithm for single cell RNA-sequencing and spatial transcriptomics

**DOI:** 10.1101/2022.05.24.493189

**Authors:** Hadas Biran, Tamar Hashimshony, Yael Mandel-Gutfreund, Zohar Yakhini

**Affiliations:** Computer Science Department, Technion - Israel Institute of Technology, Haifa, Israel; Faculty of Biology, Technion - Israel Institute of Technology, Haifa, Israel; Arazi School of Computer Science, Interdisciplinary Center, Herzliya, Israel

## Abstract

Gene expression data is complex and may hold information regarding multiple biological processes at once. We present SPIRAL, an algorithm that uses a Gaussian statistical model to produce a comprehensive overview of a plurality of significant processes detected in single cell RNA-seq or spatial transcriptomics data. SPIRAL identifies biological processes by finding sub-matrices that consist of the subset of genes involved and the subset of cells or spots. We describe the algorithmic method, the analysis pipeline and several example results. SPIRAL is available at https://spiral.technion.ac.il/.

## 2 Background

Biological processes are often characterized by changes in the levels of expression of a set of genes. The current common pipeline to discover gene expression dynamics in single cell RNA-seq (scRNA-seq) data is first to cluster the cells into cell types or states, then to perform differential expression analysis between clusters [1, 2]. Another approach is trajectory inference: a method that aims to order the cells based on a similarity measure, attempting to reconstruct the primary biological process in the data. Then, it is possible to use regression to identify genes whose expression changes along the trajectory [1].

However, both clustering and trajectory inference are usually done after dimensionality reduction, which is either performed with principal component analysis (PCA), diffusion maps or simply by filtering for highly variable genes [1–5]. While this step ensures that the principal cell partitioning or trajectory in the data would be captured, it lowers the chance of discovering less dominant biological processes. For example, processes that only involve a small fraction of the cells. Small populations of cells may also be missed [2]. Even if the clustering or trajectory inference are done on the original space (i.e. using all genes), gene expression levels are only compared between predetermined clusters or along a predetermined trajectory, making it impossible to find biological processes that do not agree with these orderings. Finally, since both clustering and trajectory inference are ultimately determined by gene expression levels, the significance of differential expression of genes between clusters or along a trajectory cannot be evaluated without correcting for this selection bias [1, 6].

Another analysis approach is to detect co-expression modules by clustering the genes. Since the clustering is done based on the gene expression values across all cells, genes which are only coexpressed in a subpopulation of cells (for example, in certain states or cell types) might not be grouped together, and structures will go undetected. Also, while gene-clustering ensures that every gene would be assigned to a cluster, it does not provide any confidence score as to the level of fit of every gene to its cluster. As a result, we might get large and non-accurate clusters of genes. The option of fuzzy clustering of genes would provide membership grades, however, they would still rely on the entire expression pattern (rather than the expression levels in an unknown relevant subset of cells). Another caveat to both cell-clustering and gene-clustering (hard and fuzzy), is that the user must specify the clustering level in advance.

Recently, spatial transcriptomics emerged, enabling the measurement of gene expression in spots laid out on a slice of tissue, maintaining both transcriptomic and spatial information[7–9]. Some analysis methods use the spatial coordinates to find spatially variable (sometimes called: spatially coherent) genes. These are genes whose expression profile fits a spatial pattern. These methods include SpatialDE[10], trendsceek[11], SPARK[12], SPARK-X[13], the BinSpect method (which is implemented in the Giotto platform)[14], GPcounts[15] and singleCellHaystack[2] (the latter can also be implemented on scRNA-seq data using its tSNE coordinates). However, all of these methods focus on identifying individual genes, and only some of them eventually cluster these genes into gene sets. singleCellHaystack does so with kmeans, SpatialDE clusters the genes based on a spatial clustering model and Giotto clusters them with hierarchical clustering. While this approach produces spatially coherent modules, it limits the flexibility of these modules. First, only the genes that were found to be spatially variable are considered. Secondly, the known caveats of gene-clustering (described above) apply here, namely that the clusters do not have significance scores and that the chance to find partly-cooperating genes is low. And third, a gene cannot be a member of more than one module. Finally, in all three methods the user must define both the number of modules, and either a p-value threshold or a number of top genes to participate in the clustering process. Given the wide scale of gene expression datasets (sizes, sequencing technologies and spatial patterns), these parameters cannot be easily guessed.

## 3 Results and Discussion

We present SPIRAL: Significant Process InfeRence ALgorithm. To our knowledge, this is the first analysis method that uses trends of gene expression as its sole detection criterion. SPIRAL detects structures: sets of genes with a similar expression pattern in a subset of cells or spots. Each of these structures reveals information that pertains to a (biological) process in the data. SPIRAL provides a list of genes, and a partitioning of the cells or spots into layers based on the expression values of these genes. The resulted list of structures provides an overview of the data, with statistical significance components, and is highly flexible: each gene and cell or spot may participate in zero, one or more of the structures.

Briefly, SPIRAL first performs fine agglomerative clustering of the cells or spots based on their full gene expression profile. Each cluster is represented by its average expression profile and referred to as “repcell”. The default number of repcells is ~ 100. This step reduces both technical noise and computational load. SPIRAL then computes the difference in expression values between every pair of repcells, for all genes. Then, in an iterative manner, it finds a gene set and a repcell-pair set in which the genes have significantly large expression-difference values (Fig. 1). The gene set and the set of repcells constitute a consistent structure, representing a consistent change in the biology of the cells or spots.

**Figure 1:**
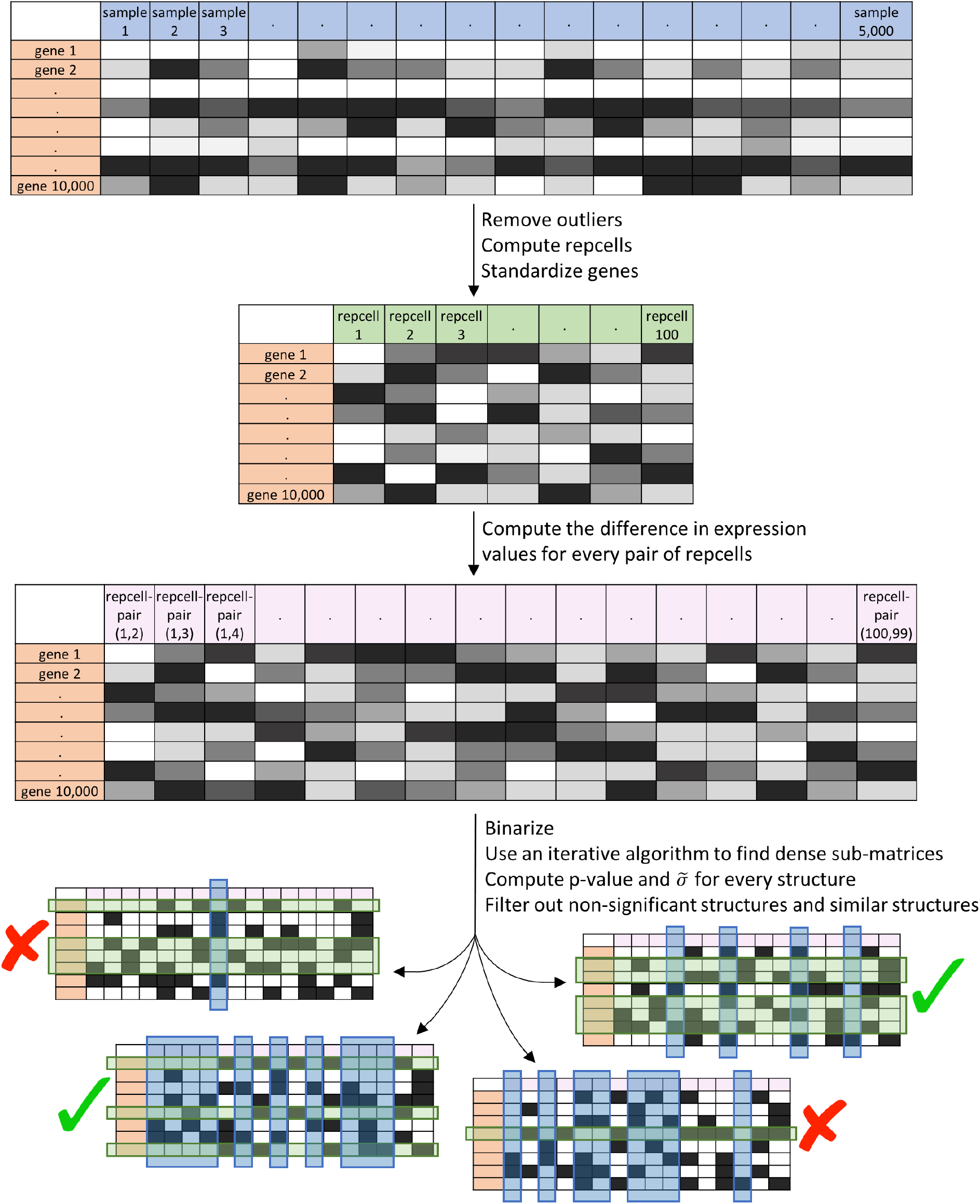
The SPIRAL algorithm pipeline. Structures marked with an X are filtered out; structures marked with a V are included in the reported structures. See text and Methods for details on the algorithm steps and the filtering criteria.

SPIRAL structures are evaluated using a statistical significance criterion based on the gene expression level differences (p-value), and a biological significance score (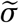- see Methods). These are used to filter out non-significant structures. Also, the list of structures is narrowed down to structures that are unique in terms of their Jaccard index of gene sets and repcell-pair sets. SPIRAL can also be applied to bulk RNA-seq. SPIRAL outputs several different visual representations of each structure, including information about the enriched GO terms of the gene set (Methods).

SPIRAL thus takes single cell or spatial transcriptomics data as input and provides significant and consistent expression patterns as output.

We applied SPIRAL to two scRNA-seq datasets (Extended Data Fig.1). The first one is of lymphoblastoid cells [16]. Fig. 2a depicts two SPIRAL structures for this dataset. SPIRAL’s assignment of cells to layers (network and left UMAP layout, for each of the structures) remarkably fit the average expression levels of the structure genes (right UMAP layout). Also, the GO enrichment analysis of the gene sets reveals that structure A captured a translational process, while structure B identified an immune process. The second dataset is of Zebrafish differentiation, taken in 7 time points post fertilization[17]. Four SPIRAL structures for this dataset are depicted in Extended Data Fig.2, revealing an upregulation in RNA splicing on hours 4 – 8 post fertilization, followed by an upregulation in ribosomal process on hours 6 – 18, in chordate embryonic development and in eryhthrocyte differentiation on hours 14 – 24 and in active cytoskeleton organization at the very late stages.

**Figure 2:**
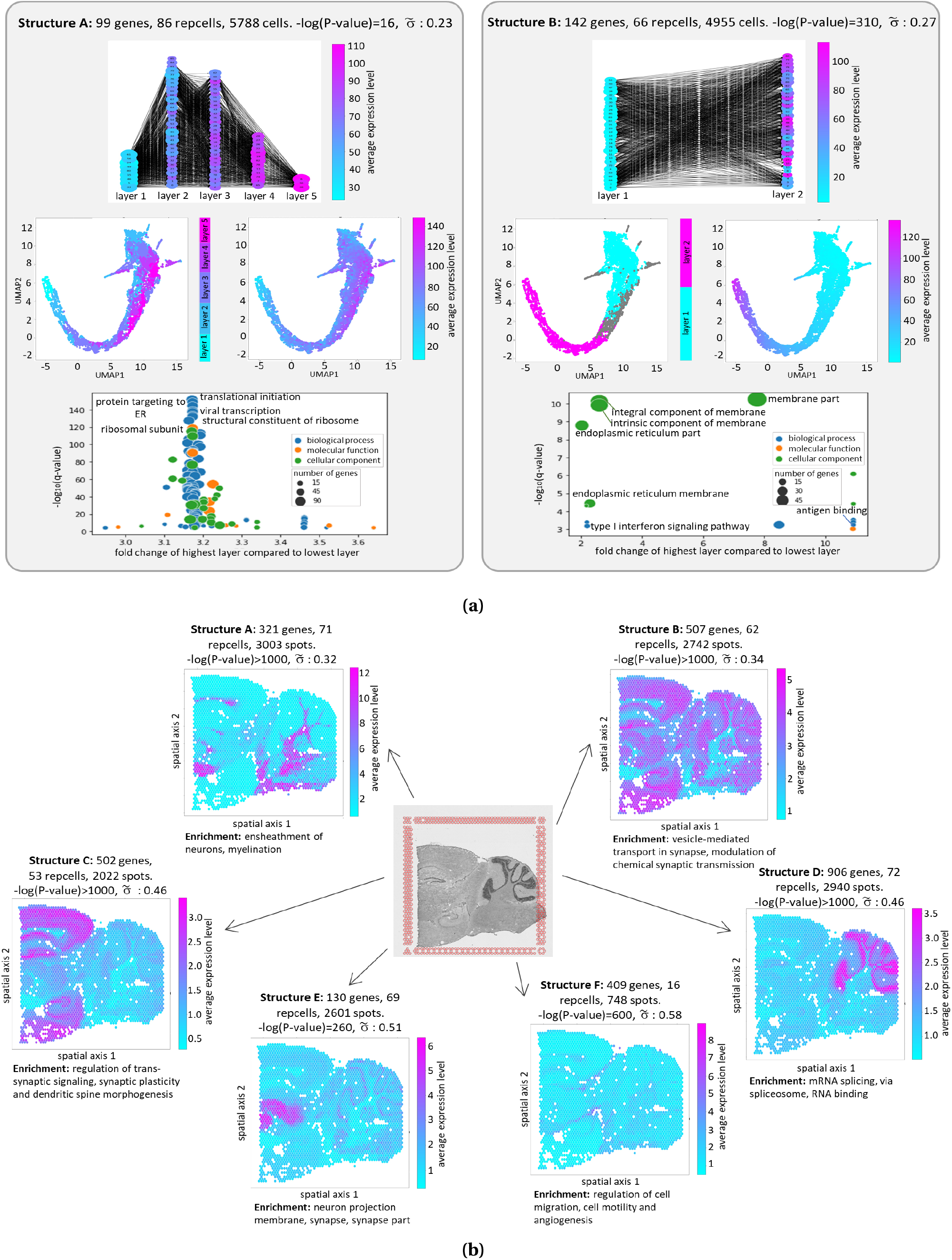
A demonstration of SPIRAL output for two datasets. **(a)** Two structures identified in a scRNA-seq dataset of lymphoblastoid cells [16]. The structures are presented in a network layout (top), in which nodes represent repcells, edges represent repcell-pairs, size of nodes corresponds to their degree; UMAP layouts (middle) and a GO term enrichment figure (bottom). **(b)** Spatial layouts of six structures identified in a spatial transcriptomics dataset of a mouse brain [18]. Average expression level refers to the average expression level of the structure genes Γ (not all genes).

We also applied SPIRAL to spatial transcriptomics data (Extended Data Fig.1). The first dataset is a sagittal-posterior section of a mouse brain from the Visium database [18]. SPIRAL detected several significant structures, spatially mapping active biological processes in the slice. For example: ensheathment of neurons and myelination, dendritic spine morphogenesis, and cell motility and angiogenesis (Fig. 2b). We note that although some angiogenesis related genes were found to be up-regulated in some clusters in Visium’s analysis of this data set [19], none of theses fit the blood vessel pattern clearly seen on structure F. The second dataset is a slice of a FFPE tissue from a normal human prostate from the Visium database [20]. SPIRAL identified and spatially mapped significant biological processes in the slice, including muscle contraction, angiogenesis and cellular response to chemical stimulus (Extended Data Fig.3).

We applied SPIRAL to two bulk RNA-seq datasets (Extended Data Fig.1). The first consists of 31 bulk samples representing undirected differentiation of human embryonic stem cells, taken at 16 time points on days 0 and 7 – 21 of differentiation (Methods). SPIRAL identified upregulated translational activity at hours 0 – 10 and anatomical structure development activity at hours 12 – 18. Regulation of cell differentiation, cell migration and T-cell activation were upregulated at hours 15 – 21. SPIRAL also detected a batch effect between replicates (Extended Data Fig.4). The second dataset covers 50 samples of mouse B cells treated with anti-IgM mAb to mimic B cell receptor stimulation, with expression measured every 15 minutes during six hours post stimulation [21, 22]. SPIRAL identified increased regulation of B cell activation at 0 – 30 minutes post stimulation, increased regulation of cell morphogenesis and cell differentiation at 15 – 90 minutes post stimulation, increased lymphocyte and leukocyte differentiation at 45 – 225 minutes, and increased translational activity at 165 – 360 minutes. SPIRAL also detected batch effects (Extended Data Fig.5).

## 4 Conclusions

The computational biology community has developed a wide variety of analysis techniques and algorithms for scRNA-seq and spatial transcriptomics data[**bacher2016design**, **zeng2022statistical**]. However, most of these methods are aimed at finding the main data partition or time-line. In this paper we presented methods that are highly flexible and fully adjustable to the data. Our approach captures all data features and allows for the determination of statistically significant structures, distinguished from noise. More specifically, SPIRAL detects structures combining selection on both gene and sample axes. SPIRAL uses a Gauss based statistical approach for assessing significance. SPIRAL is available at https://spiral.technion.ac.il/.

## 5 Methods

### 5.1 Related computational problems

Our work is inspired by the work of Ben-Dor *et al*. [23], who developed a greedy algorithm that searches for order-preserving submatrices (OPSMs) in gene expression matrices. OPSMs are defined as submatrices in which it is possible to find a permutation of the samples (columns), such that the expression values of every gene (row) in the sub-matrix monotonically increase in respect to that permutation [23, 24]. OPSM’s main drawback is its insistence on a perfectly increasing set of expression values for all involved genes, which may be too strict for biological data. Two research groups[25, 26] presented “relaxed” versions of the problem, both by using ideas from sequential pattern mining, and specifically-using a depth-first traversal of a prefix tree to find frequent subsequences. However, these approaches also lack statistical considerations in defining the required properties of the structures of interest.

#### Searching dense sub-matrices in a binary matrix

A central step of the SPIRAL protocol is the identification of dense submatrices in a large binary matrix, where we define “dense” as having at least a minimum fraction of 1’s (denoted by *μ*) in every column and in every row of the submatrix. We note that methods that aim to find “all 1’s” sub-matrices in binary matrices are irrelevant due to the noisy character of biological data, and specifically scRNA-seq data. So, we focus here on the problem of finding dense sub-matrices in binary matrices.

In the past two decades, several studies have proposed heuristic algorithms to solve similar problems. Both Koyuturk *et al*. [27] and Uitert *et al*. [28] focused on finding submatrices with a significantly large fraction of 1’s. We note that this definition of the problem is *global*, that is, it requires density in the entire sub-matrix and not in individual columns or rows. Their proposed heuristic algorithms begin with a random set of columns or rows, and then iteratively improve the choice of row and column sets, until no further improvement in the significance score is possible. In both cases, the significance scores are computed based on Chernoff bounds and the null assumption is that the binary matrix was generated by a Bernoulli distribution on each of the matrix elements, with success probability 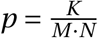 (where the input matrix has *M* rows by *N* columns with overall *K* values of 1). The independence assumption does not apply to our model (see Section 5.2.5). Our definition of a dense sub-matrix is also slightly different (see Definition 1). Nonetheless, our algorithm also uses an iterative approach, in the spirit of Koyuturk *et al*. [27] and Uitert *et al*. [28].

The problem of finding dense sub-matrices in a binary matrix can also be formulated as the quasi-biclique problem. Given a binary matrix, let us construct a bipartite graph such that the vertices on the left correspond to the matrix rows, the vertices on the right correspond to its columns, and an edge exists between vertex *i* on the left and vertex *j* on the right iff element *i*, *j* of the binary matrix equals 1. Then, the problem of finding a significantly dense sub-matrix in the binary matrix is equivalent to the problem of finding a set of vertices on the left and a set of vertices on the right of the bipartite graph such that the number of edges between the first set and the second set is significantly high.

For a bipartite graph *G* = 〈*V*_1_, *V*_2_, *E*〉 with two disjoint sets of vertices *V*_1_, *V*_2_ and with edges *E*, Mishra *et al*. [29] define an *ϵ-biclique* as two sets of vertices 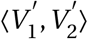 such that 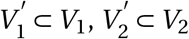, and every vertex in 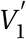 is connected to at least 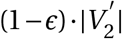 vertices in 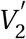 (where *ϵ* ∈ (0, 1]). This definition is *local* but *asymmetric*. That is, in the original problem it applies to every row but not to every column in the sub-matrix. Their algorithm for finding such structures relies on exhaustive sampling.

Li *et al*. [30] define a *δ%-tolerance quasi-biclique subgraph* as two sets of vertices 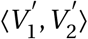 such that 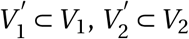, and every vertex 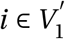 is disconnected from up to 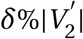 vertices in 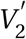, and vice versa. This definition is *local* and *symmetric*, so it applies separately to every column and row in the original problem. The authors state that for a predefined set of vertices, it is possible to perform iterations to find a *δ*%-tolerance quasi-biclique that contains that set. However, no explicit algorithm for this problem is described.

We also chose to define our structures of interest locally and symmetrically, to avoid skewness of the quasi-biclique [30], which translates to rows or columns that are loosely related to the submatrix. In fact, our definition of a *μ-dense sub-matrix* (see Definition 1) is equivalent to Li *et al*. [30]’s *δ%-tolerance quasi-biclique subgraph* for 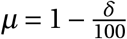.

Our work extends the above literature by:

- We propose a different probabilistic modelling approach for dense submatrices, that is more adequate to matrices wherein each row is not assumed to represent a distribution with a diagonal covariance matrix. That is: elements within rows can be statistically dependent. This more general approach is necessary for modelling progression in scRNA-seq data. Our approach specifically applies to rows with any multivariate Gaussian distribution.
- We use our approach to find significant trends in matrices representing scRNA-seq as well as spatial transcriptomics data. To do so, we model such data as coming from Gaussian distributions and then translate trends to structures in a binary matrix derived from this representation.
- We assign a significance score to the trends discovered as above and further investigate the related biology.
- We develop a simple and efficient algorithmic approach to all components of the above discovery process.

### 5.2 The SPIRAL protocol

We start by observing an expression matrix *A* with *M* rows (genes) and *N* columns (cells for scRNA-seq or spots for spatial transcriptomics). We assume that there is a set of genes acting in concordance with one another to execute some biological process on a set of cells. Some of these genes may be driving the process and some associated with it in other ways. The coordinated expression pattern can be inferred from the data, providing an indication of the biological process.

We aim to find this process by finding a collection of pairs of columns (cells\spots) in which the expression of the relevant genes changes significantly and concordantly.

#### 5.2.1 Pre-processing the data

To eliminate the effect of outliers and technical artifacts, we filter out cells\spots (scRNA-seq\spatial transcriptomics respectively) with abnormally high or low number of expressed genes (which may be doublets and empty cells) [1]. We then identify mitochondrial genes by querying the Ensembl BioMart database [31], and remove cells\spots with a large percent of mitochondrial RNA, as this could suggest cell stress [32, 33]. As a final pre-processing step, we normalize each cell\spot to the median number of counts per cell\spot.

#### 5.2.2 Computing representing cells

In order to avoid dropouts and inaccuracies which stem from technical issues, we cluster the cells\spots into small low variance clusters. Then, for each cluster, we consider its member cells\spots to be biological replicates and compute their average expression profile. This average expression profile is the “representing cell” of that cluster. This idea is inspired by Iacono *et al*. [34]’ iCells. Specifically, the clustering is executed here by employing hierarchical agglomerative clustering with the ’ward’ linkage. The number of clusters is set to 100 for datasets with 3000 cells\spots or more, and to the number of cells\spots divided by 30 otherwise. After clustering, clusters with less than 10 cells\spots are discarded. For convenience, we refer to the representing cells as “repcells”.

The expression profiles of all repcells are then stored in a new expression matrix *A_c_* with size *M* by *N_c_*, whose rows are genes and columns are the repcells.

This step improves accuracy by averaging biological replicates, eliminates dropouts, and reduces the number of samples while preserving all genes. The reduction in data size shortens the execution time of the SPIRAL algorithm without removing significant information from the data (as demonstrated on a synthetic dataset in Section 5.4).

#### 5.2.3 Registering changes in expression values

We first standardize each gene separately. In this step, the (*g*, *j*)th entry in the normalized matrix 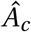 is computed as 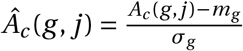, where *m_g_* is the gene average expression and *σ_g_* is its standard deviation over all *N_c_* repcells. Explicitly: 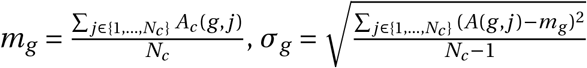.

We compute the difference in expression values between every two repcells, for all genes, by multiplying 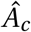 on the right by a pairwise subtraction matrix *B*:

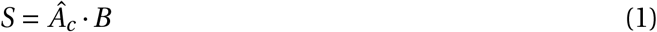

*B* is constructed so that all repcell pairs (in both directions) will be evaluated. It has *N_c_* rows and *N_c_* · (*N_c_* – 1) columns. For example, for 4 repcells (*N_c_* = 4), *B* would be:

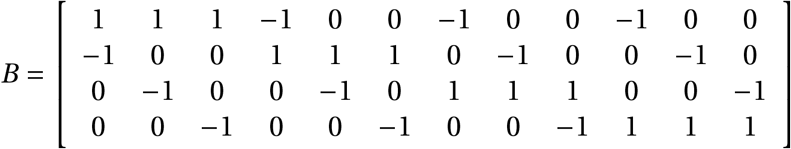

The resulting matrix *S* represents the repcells’ differences. It has rows that correspond to genes, and columns that correspond to ordered repcell-pairs. An entry *S*(*g*, (*i*, *j*)) is the difference in expression levels of gene *g* between a repcell *i* and a repcell *j*, i.e:

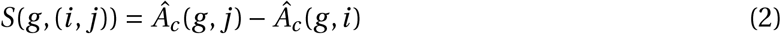

#### 5.2.4 Seeking dense sub-matrices

We now wish to find a set of genes (rows in *S*) and a set of repcell-pairs (columns in *S*) that would constitute a sub-matrix that is populated by almost exclusively high values (a formal definition is given later in this section). For this purpose we first binarize the matrix *S* such that

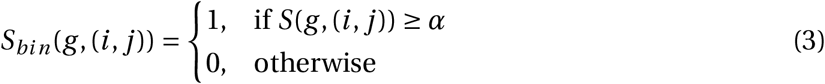

where *α* is a parameter determining the minimal number of standard deviations (of a gene) to be considered a significant change in expression (since we can also write: 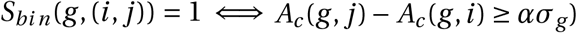.

We now wish to find a dense sub-matrix in the binary matrix *S_bin_*. For that purpose, we introduce the following definition:

##### Definition 1

*a μ**-dense sub-matrix** is a collection of rows* Γ *and columns* Π *of a binary matrix such that every row in* Γ *has at least μ*|Π| *non-zero values within the columns* Π *and every column in* Π *has at least μ*|Γ| *non-zero values within the rows* Γ.

Formally: *S_bin_* (Γ, Π) is *μ-dense* if:

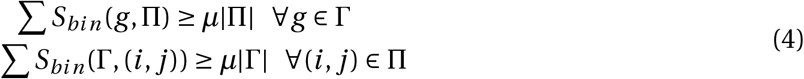

where *μ* is a density parameter in the range (0, 1], which, in practice, is typically close to 1.

##### The iterative dense sub-matrix algorithm

Inspired by both Koyuturk *et al*. [27] and Uitert *et al*. [28], we aim to find *μ*-dense sub-matrices by using an iterative algorithm (see Algorithm 1). The algorithm initiates each of its *T* iterations with a random path of repcells with length *L*, and then repeatedly improves its choice of genes (rows) and repcell-pairs (columns), until convergence to a single structure (when the structure remains unchanged after executing the while loop). All the results in this paper were obtained with *L* = 3 and *T* = 10000.

###### Algorithm 1 The iterative dense sub-matrix algorithm

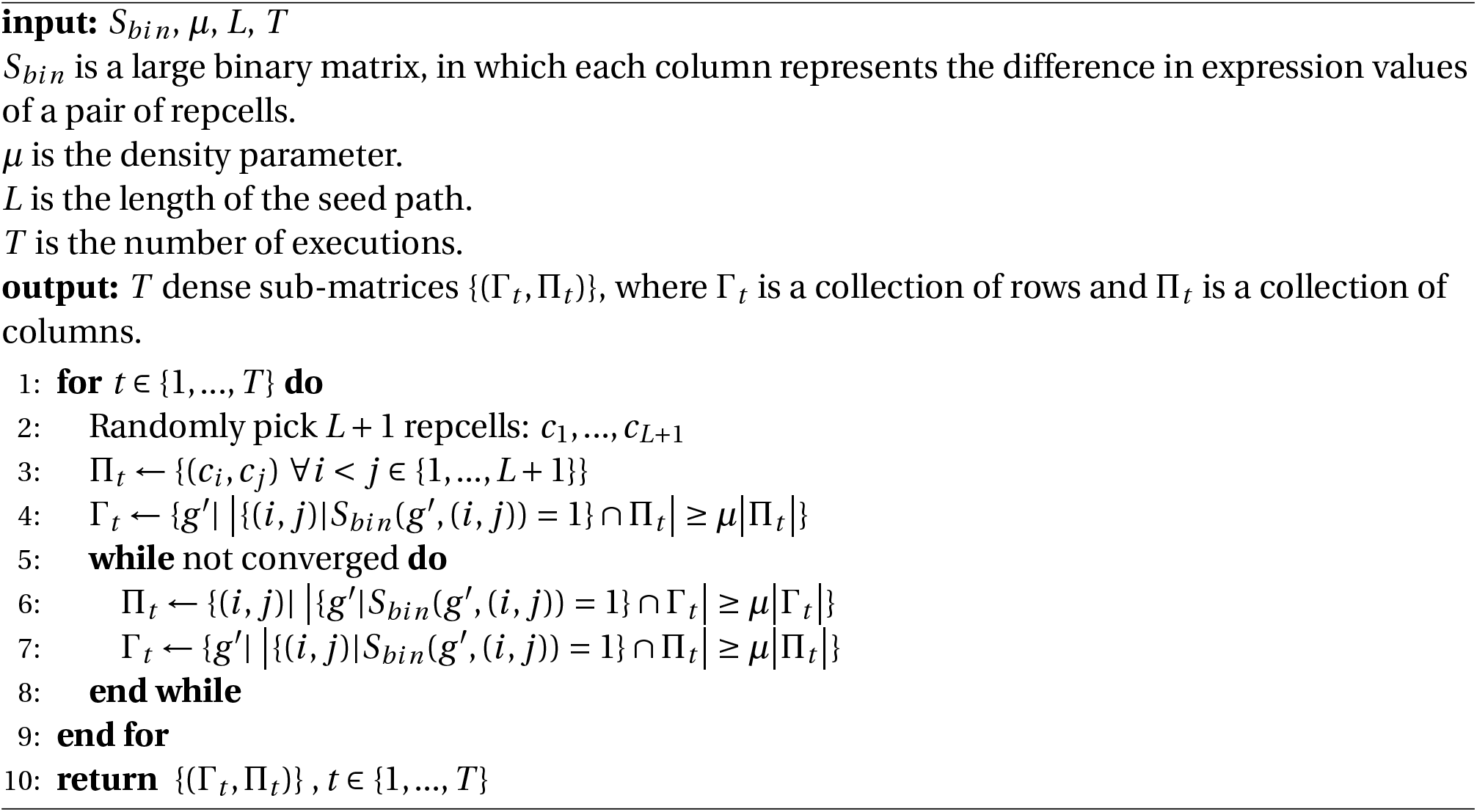

#### 5.2.5 Evaluating the statistical significance of a given sub-matrix

We now wish to calculate a score representing the statistical significance of a *μ*-dense sub-matrix of the matrix *S*, which consists of the set of columns (repcell-pairs) Π and the set of rows (genes) Γ. We clarify that although the sub-matrix was found using a binarized version of *S*, the significance evaluation is done on the original *S* which contains continuous values.

This sub-matrix in *S* was originally produced by

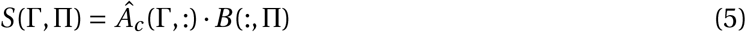

##### Computing the statistical significance of a single gene and a set of repcell-pairs

For the null model, we assume that a standardized expression value of a gene *g* in a repcell *i* is distributed as 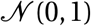. Moreover, we assume independence and thus the set of standardized expression values ***X_g_*** of a gene *g* in all *N_c_* repcells is distributed as

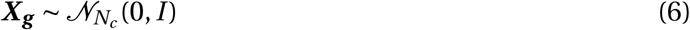

and any row *g* of 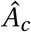 is, under the null model, a single instance drawn from that distribution.

For a set of repcell-pairs Π and a single gene *g*, we now define 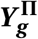 to be an affine transformation of ***X_g_*** (both 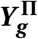 and 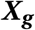 are row vectors):

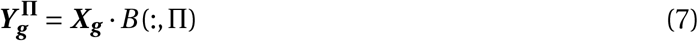

In the null case, the set of values *S* (*g*, Π) is a single draw of 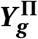.

At this point we wish to clarify that although the standardized expression values of a gene *g* (denoted by the vector ***X_g_***) are assumed, under the null model, to be collectively independent, such assumption regarding their pairwise differences (represented by 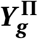) is not valid. Also, if *B* (:, Π) is not full rank, 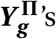 distribution is singular.

We now define:

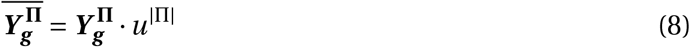

where *u*^|Π|^ is a column vector with all elements equal to 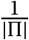, i.e. an averaging vector.

We can conclude from equations (6), (7) and (8) that 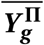 is a real valued normal variable with mean 0 and standard deviation 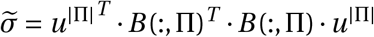. That is,

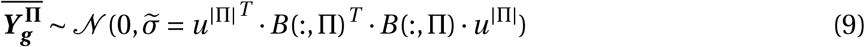

(see Figure 3).

**Figure 3:**
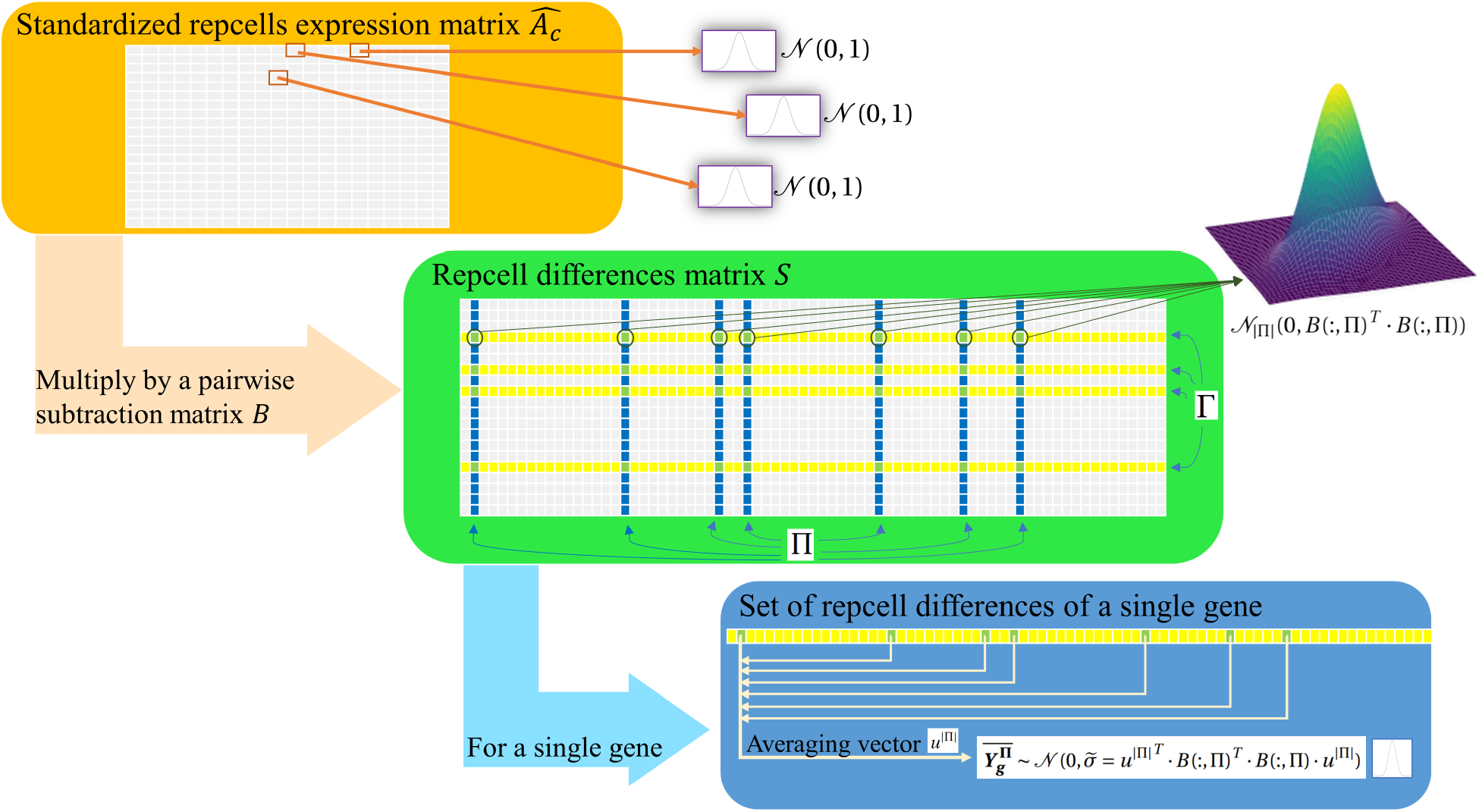
A flow chart of the null assumption of our model.

Therefore, the significance of the observed average change in expression values of gene *g* over the set of repcell-pairs Π can be computed as

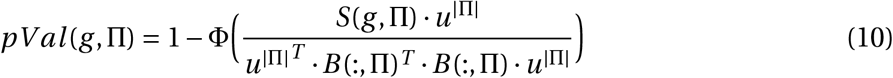

where Φ is the cumulative distribution function of the standard normal distribution, the numerator is the observed average of values in *S*(*g*, Π) (the average of the observed expression differences of *g* throughout Π), and the denominator is 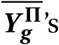 standard deviation, as developed above.

##### Computing the significance of a set of genes and a set of repcell-pairs

For the null model we also assume that the genes are collectively independent. We can then compute the significance of a given sub-matrix with columns (repcell-pairs) Π and rows (genes) Γ by multiplying the significance scores of the individual genes:

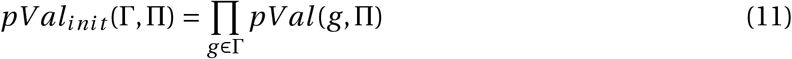

##### Correcting for multiple testing

We correct the initial p-value for multiple testing as follows

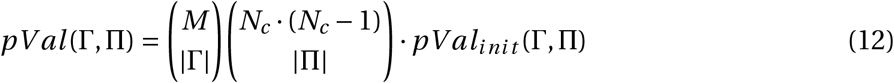

where *M* is the total number of genes and *N_c_* is the total number of repcells. Note that *N_c_* · (*N_c_* – 1) is then the total number of repcell-pairs.

#### 5.2.6 Evaluating the structure of repcells consisting of the sub-matrix columns

The significance score indicates how statistically rare a discovered sub-matrix would be under the null hypothesis. However, we argue that due to the properties of real-life data, there is a need for a second measure to asses biological significance. For that purpose, we examine the structure consisted of the set of repcell-pairs.

We may think of a set of repcell-pairs Π = {(*i*_1_, *j*_5_), (*i*_3_, *j*_8_), …} as a network in which the nodes are repcells, and an arrow is drawn from repcell *i* to repcell *j* only if the repcell-pair (*i*, *j*) is in Π.

Using three toy examples, we will now demonstrate that different sub-matrices may have similar significance scores (and also, similar number of repcell-pairs), but very different contributions to the understanding of biological processes in the data. We will then offer a second measure that addresses biological significance.

Consider the structures in Figure 4: All three have 33 repcell-pairs (arrows), and their numbers of participating genes are 100, 30 and 50 for structures (a),(b),(c) respectively. We assume *α* = 1, *μ* = 1, and that the difference in expression values along each of the arrows in structures (a) and (b) is exactly 1 for each of the genes (i.e. *S* (*g*, (*i*, *j*)) = 1 ∀*g* ∈ Γ, (*i*, *j*) ∈ Π), and for structure (c) this is also true for all arrows but the ones connecting the first and the last layers (i.e. *S*(*g*, (*i*, *j*)) = 1 *if* [(*i* ∈ {0, 1, 2} & *j* ∈ {3, 4, 5, 6}) || (*i* ∈ {3, 4, 5, 6} & *j* ∈ {7, 8, 9})], otherwise *S*(*g*,(*i*, *j*)) = 2). In this case their initial significance scores (computed as in equation (11)) are 1*e*-79.0, 1*e*-68.5 and 1*e*-64.4 respectively. Despite the similarity in significance scores, it is clear that while structures (b) and (c) might suggest evidence to a biological process in the data, structure (a) might as well be a result of a technical issue involving one outlier.

**Figure 4:**
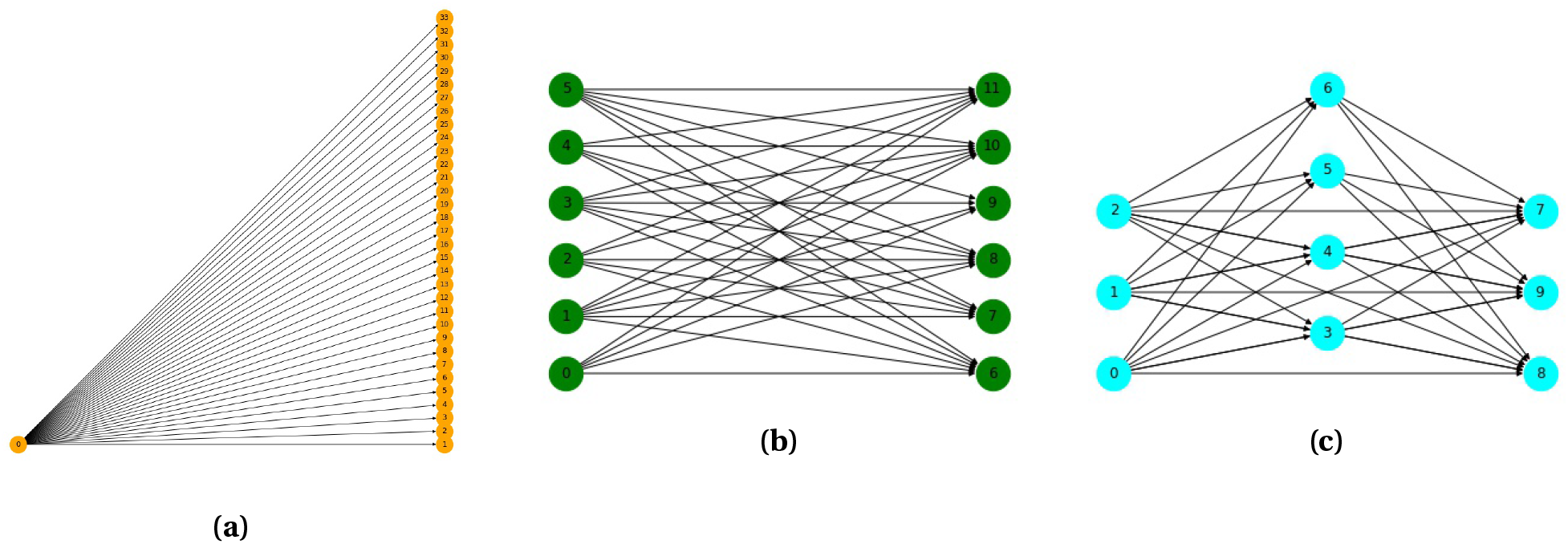
Toy examples of structures.

As mentioned before (see equation (9)), the average of differences along all repcell-pairs in the structure is distributed in the null case as a normal variable. Consider the standard deviation of this variable, denoted by 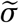. When it is lower, the structure is more complex and might offer more biological insights. For structures (a), (b) and (c), 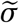 is 1.02, 0.58 and 0.52 respectively, allowing us to easily determine which of the structures is more informative.

#### 5.2.7 Filtering sub-matrices

After evaluating the statistical significance of the resulting sub-matrices, we filter out non-significant ones (p-value above 1*e*-15). Note that in random data, we get no structures with p-value below this threshold. Then, we perform a disjointification-like process to avoid having very similar sub-matrices in the algorithm output list. The process begins by sorting the sub-matrices in an ascending order of their structures’ 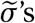 (see Section 5.2.6). Then, we go through the list of submatrices and remove every sub-matrix that is similar to any of the former sub-matrices in the list, where two sub-matrices are considered similar if the Jaccard index of their gene lists is above 0.75 or if the Jaccard index of their repcell-pair lists is above 0.5. At this point, different datasets may result in very different numbers of structures. To assure that the resulted number of structures in within a certain range, we cut the structure list (still, sorted by structures’ 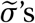) at 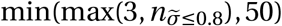, where 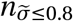 is the number of structures with 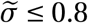. We note that in the web tool, a user may narrow down this list by adjusting the Jaccard index threshold of the gene lists.

### 5.3 SPIRAL visualizations and gene enrichment

#### 5.3.1 Producing the network layout and partitioning the repcells to layers

For structures that can be drawn as a bipartite graph (such as the toy examples in Figs. 4a and 4b), we can label the repcells on the left (layer 1 in the network) as “low expression” and the repcells on the right (layer 2) as “high expression”. However, for other structures, there are several options of partitioning into network layers. We chose to draw the network from right to left, so that the arrows are short as possible (see Algorithm 2). After the partition is established, we assume that in general the expression levels of the structure genes are increasing from left to right: the repcells on layer *i* would likely have lower expression values of the structure genes than the repcells on layer *i* + 1. In the simple network visualization, the repcells are colored based on their assigned layer (see Figure 6a). However, to test the former assumption, SPIRAL also produces a network visualization in which the repcells are colored based on their average expression level of the structure genes (see Figure 6b). The sizes of nodes in the network layout correspond to their degree: the number of edges connecting them to other repcells. This is a visual indication of the level of connectivity of each repcell in the structure.

#### 5.3.2 Visualizing significant sub-matrices on a PCA\UMAP layout of repcells

SPIRAL visualizes the structure over a PCA \UMAP layout by coloring the repcells based on their assigned layers (see Section 5.3.1 and Figure 6c). As with the network layout, SPIRAL also produces a more informative visualization by coloring the repcells based on their average expression level of the structure genes (see figure 6d).

##### Algorithm 2 The network drawing process

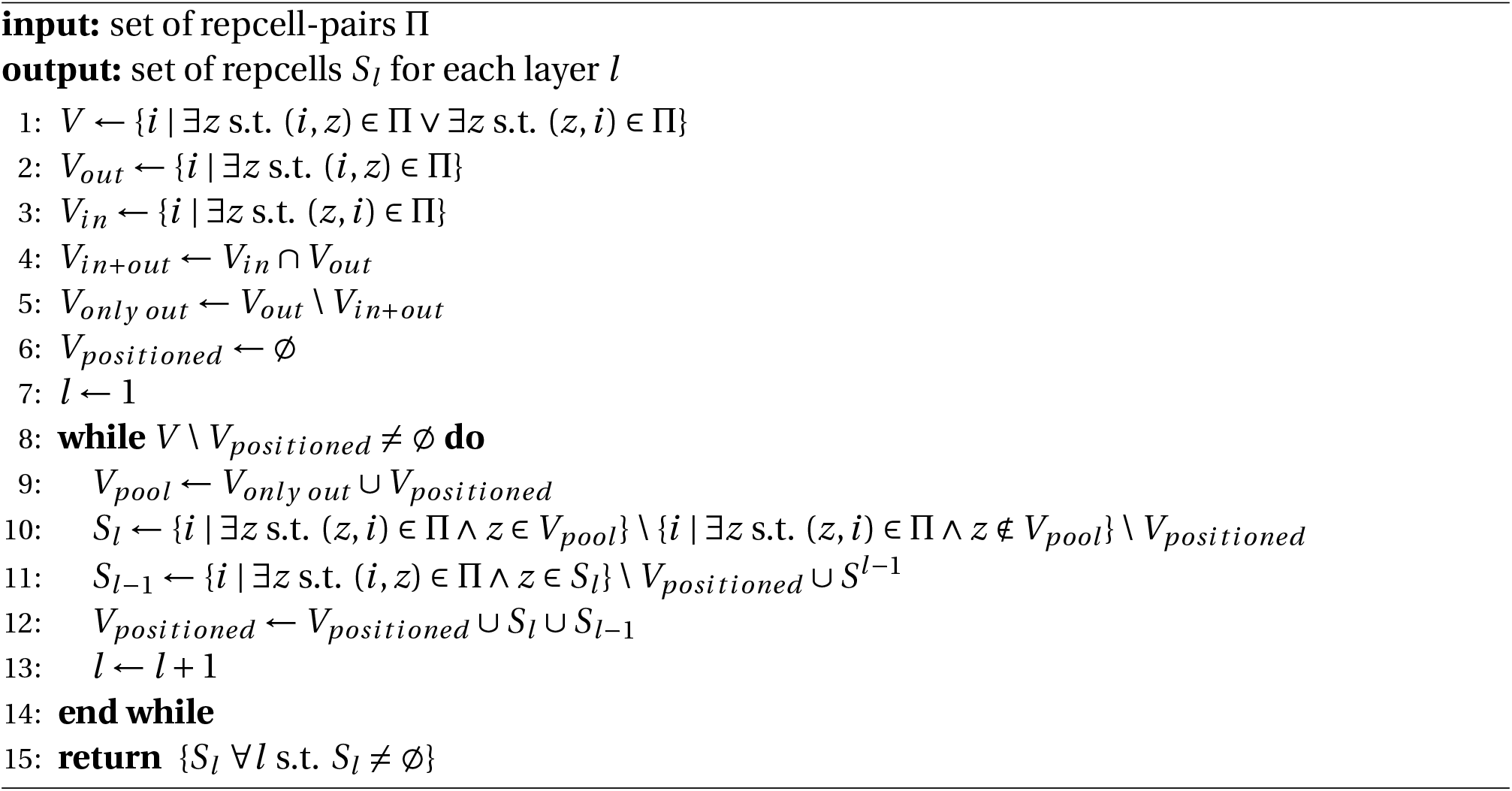

#### 5.3.3 Visualizing significant sub-matrices on a PCA\UMAP\spatial layout of cells\spots

SPIRAL visualizes structures on a PCA\UMAP layout of the original cells\spots, or on the spatial coordinates of the spots, by assigning each cell\spot with a layer based on its repcell (see 5.3.1 and Figure 6e). SPIRAL also offers a similar visualization, in which the assigned layers are ignored, and instead-the cells\spots are colored based on their average expression level of the structure genes (see Figure 6f).

#### 5.3.4 Evaluating the set of genes Γ

In order to asses whether the structure genes are enriched for known biological processes or functions, SPIRAL performs a GO enrichment query through GOrilla [35, 36].

### 5.4 Validating our algorithmic approach on synthetic data

We used Splatter [37] to synthesize scRNA-seq data of 7000 cells and 10000 genes, with a branching lineage. The lineage was constructed such that each cell belongs to one of three paths (with probabilities 0.2, 0.2, 0.6 to belong to Path1, Path2, Path3 respectively). Path2 and Path3 begin where Path1 ends. Also-the location of each cell along its path (i.e. step) is also known (overall 1000, 1000, 3000 steps in Path1, Path2, Path3 respectively). We defined the differential expression factor “de.facLoc” to be 0.01, which is considered small (which translates to small differences between cells in different paths) [38]. A PCA layout of the synthetic data is presented in Figure 5.

**Figure 5:**
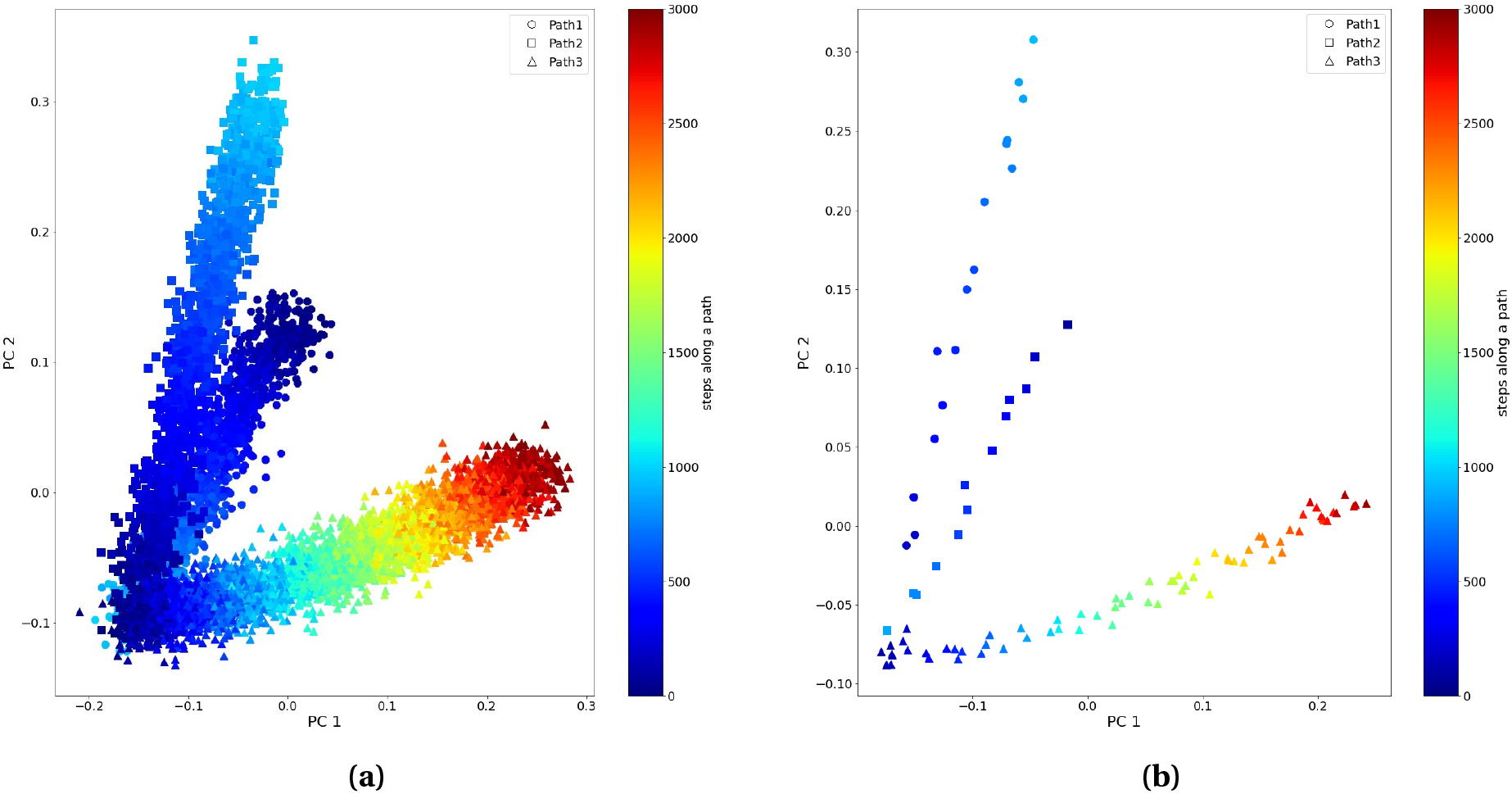
PCA layouts for Splatter synthetic data and repcells. Shapes correspond to paths, colors correspond to steps. **(a)** PCA for synthetic data. **(b)** PCA of repcells for synthetic data.

We then ran the SPIRAL algorithm on the synthetic dataset (with *α* = 0.5, *μ* = 0.95). A PCA of the 100 repcells is presented in Figure 5b. For intuitive visualization, the path of each repcell in Figure 5b was determined by applying a majority vote of the paths of the original synthetic cells that were part of the cluster of that repcell. Then, the repcell’s step was determined by averaging the steps of the “winning” cells in the path vote.

We present here seven SPIRAL structures for this dataset. For the first one, which involves 340 genes, 6666 cells and 93 repcells, we present a panel of visualization layouts in Figure 6. The network layouts in Figs. 6a and 6b provide information regarding the level of connectivity and skewness of the structure. Here all repcells (nodes) have similar size (which means their degrees are similar, see 5.3.1), so the structure is well-connected and not skew. The PCA layouts of repcells in Figs. 6c and 6d allow us to place the process on a time line, based on our preliminary information of the dataset. Here, we can conclude that the structure genes amplify their expression along the lineage of Path3. The PCA layouts of the original data in Figs. 6e and 6f ensure that the repcells faithfully represent the original data. SPIRAL exact partition to layers, depicted by the colors in the three left figures, tells us about how this structure was spot by the algorithm. However, more detailed insights regarding change over time can be captured by looking at the three right figures, in which the colors correspond to average expression level of the structure genes.

**Figure 6:**
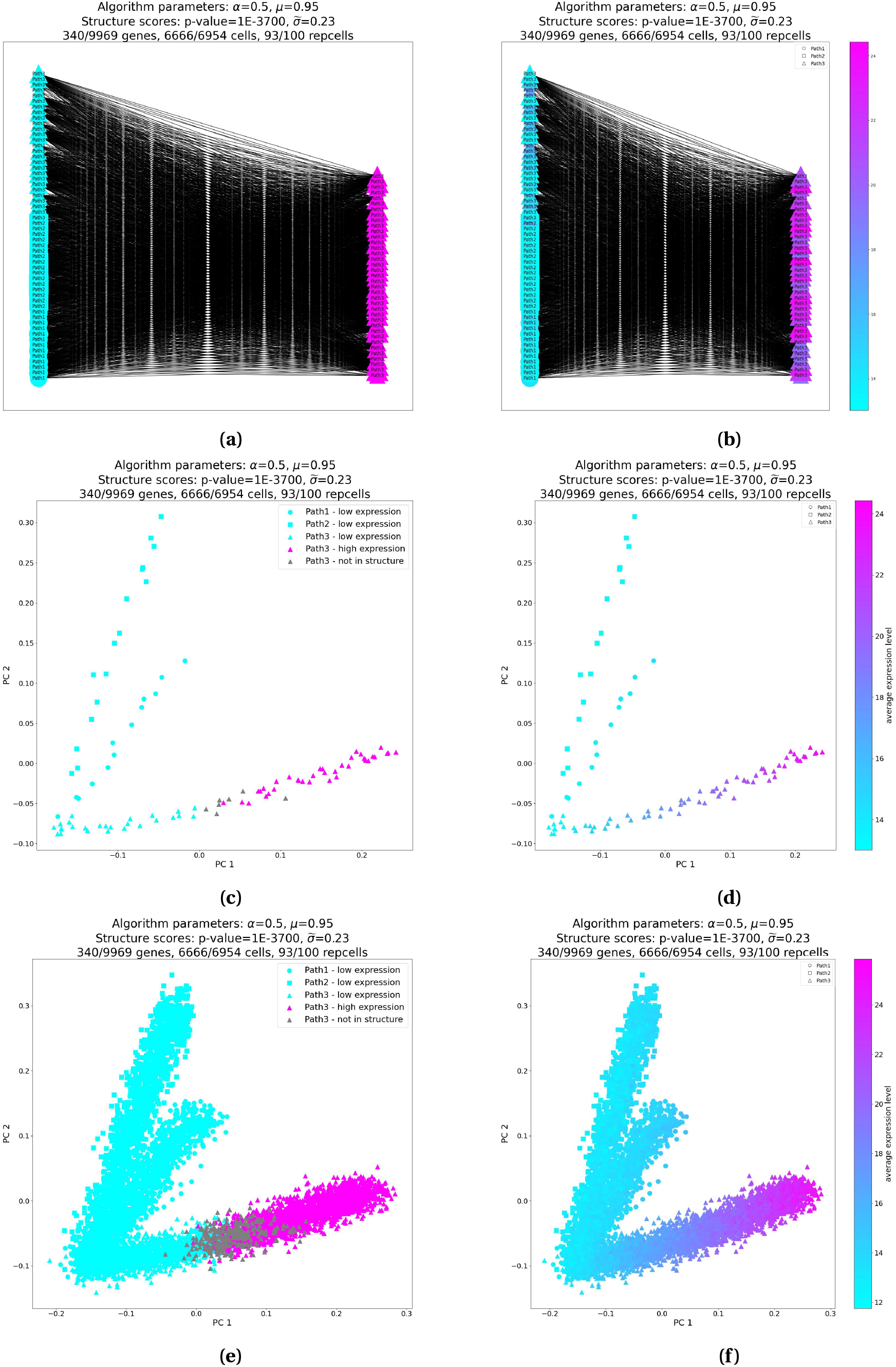
Different visualizations of a single SPIRAL structure found in synthetic data produced by Splatter. The structure consists of 340 genes that increase their expression along Path3. **(a),(b)** Network layouts. **(c),(d)** PCA layouts of the repcells. **(e),(f)** PCA layouts of the cells. **(a),(c),(e)**-colored by structure layers. **(b),(d),(f)**-colored by the average expression level of the structure genes.

The PCA layouts of additional six structures for this dataset are presented in Figure 7.

**Figure 7:**
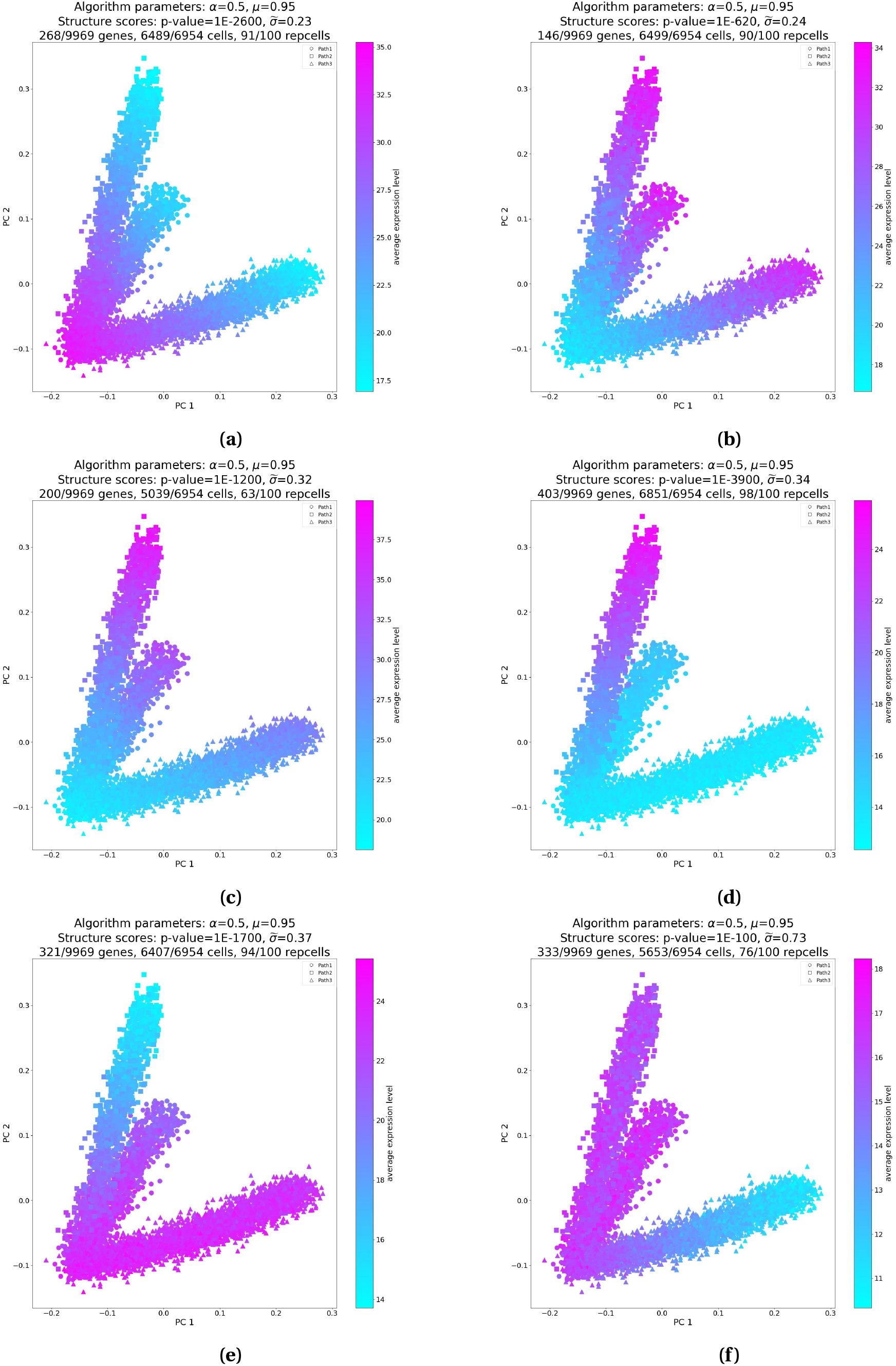
PCA layouts for structures of Splatter synthetic data. The structures indicate that: **(a)** 268 genes are amplified at the end of Path1 and then reduced again along Path2 and Path3. **(b)** 146 genes reduce their expression along Path1 and are then amplified along Path2 and Path3. **(c)** 200 genes are somewhat expressed at the beginning of Path1, then they are reduce, and then they are amplified at the end of Path2. **(d)** 403 genes that are not expressed at all throughout Path1 and Path3, are amplified along Path2. **(e)** 321 genes are only expressed in Path1 and Path3. **(f)** 333 genes are only expressed in Path1 and Path2.

### 5.5 SPIRAL web tool

SPIRAL web tool has a friendly walkthrough data upload process. When SPIRAL results are ready (typically 4-6 hours post data loading), an indication is sent by email. Then, the user can observe the structure visualizations in the result panel webpage. It is also possible to download the result table and all the relevant visualizations.

### 5.6 Bulk RNA-seq data generation

The following pipeline was used to generate the bulk RNA-seq dataset of human embryonic stem cells (hESCs) differentiation.

#### 5.6.1 Spontaneous differentiation of hESCs

Suspension cultures of TE03 cells were routinely maintained in hESC-medium with 100 ng/ml bFGF. To induce spontaneous differentiation, cells were transferred to differentiation medium consisting of DMEM/F-12 supplemented with 10% FBS (BI, Cat# 04-002-1A), 10% KnockOut Serum Replacement (KSR), 1 mM L-glutamine, 1% non-essential amino acids, and 0.1 mM *β*–mercaptoethanol, media changed every 1-2 days. After one week media was replaced with FBS media, consisting of DMEM supplemented with 20% FBS 1 mM L-glutamine, 1% non-essential amino acids, and 0.1 mM *β*–mercaptoethanol, media changed every 2-4 days for 2 weeks. Samples collected on the indicated days, RNA extracted by TRI-reagent according to protocol.

#### 5.6.2 Library preparation and sequencing

2 ng RNA of each sample was taken for library preparation using the CEL-Seq2 protocol[39]. Briefly, The CEL-Seq primer selects for polyA RNA via an anchored polyT stretch, and adds a sample specific barcode, a UMI, the Illumina adaptor and a T7 promoter sequence to the resulting dsDNA (2 different CEL-Seq primers added to each sample as technical replicates). At this point, multiple samples can be pooled together, and an In-Vitro Transcription (IVT) reaction is performed to linearly amplify the RNA. The second Illumina adaptor is introduced at a second Reverse Transcription step (RT) as an overhang to a random hexamer primer. A short PCR reaction selects for the 3’ most fragments and add the full Illumina adaptors needed for sequencing. Sample and molecule origin are identified by read 1, and gene of origin is identified by read 2. Library was sequenced on Nextseq550, 25 bases for read 1 and 60 bases for read 2.

#### 5.6.3 Bioinformatic analysis

Demultiplexing was performed using Je-Demultiplex [40] in Galaxy platform. R2 reads were split into their original samples using the CEL-Seq barcode from R1. Reads from two technical replicates were joint for further processing. The reads were cleaned using Cutadapt[41] for removal of adaptors, polyA, low-quality sequences (Phred< 20) and short reads (<30bp after trimming). Reads were mapped to the GRCh38 genome (Ensembl) using STAR package[42] to create Bam files. We used SAMtools[43] to convert BAM to SAM file and index the files. Then the reads were annotated and counted using HTseq-count package[44].

## 6 Data availability

The scRNA-seq count matrix of lymphoblastoid cells[16] was downloaded from https://www-ncbi-nlm-nih-gov/geo/query/acc.cgi?acc=GSM3044892. The Zebrafish differentiation count matrices (one for each time point) [17] were downloaded from https://www-ncbi-nlm-nih-gov/geo/query/acc.cgi?acc=GSE112294. The Feature/barcode matrix (filtered) of the sagittal-posterior section of a mouse brain, as well as the corresponding spatial coordinates and the H&E histological image of the slice (Fig. 2b) were downloaded from 10x-Genomics’s website at https://www.10xgenomics.com/resources/datasets/mouse-brain-serial-section-2-sagittal-posterior-1-standard-1-1-0. The Feature/barcode matrix (filtered) of the human prostate, as well as the corresponding spatial coordinates and the H&E histological image of the slice (Extended Data Fig.3) were downloaded from 10x-Genomics’s website at https://www.10xgenomics.com/resources/datasets/normal-human-prostate-ffpe-1-standard-1-3-0. The bulk RNA-seq count matrix of mouse B cells, treated with anti-IgM mAb[21], was downloaded from https://www.ncbi.nlm.nih.gov/geo/query/acc.cgi?acc=GSE129536. The bulk RNA-seq count matrix of human differentiation is available at https://github.com/hadasbi/SPIRAL.web.tool/blob/7b7ec5da55da7071d0fceb42a6b67d0a30019c49/human_differentiation_count_matrix.csv.

## 7 Code availability

The algorithm and website code is available at https://github.com/hadasbi/SPIRAL.web.tool.

## 8 Acknowledgements

We would like to thank Tamar Lahav and Amir Argoetti for useful discussions.

## 9 Extended data figures

### 9.1 Table 1: datasets

**Extended Data Fig.1:**
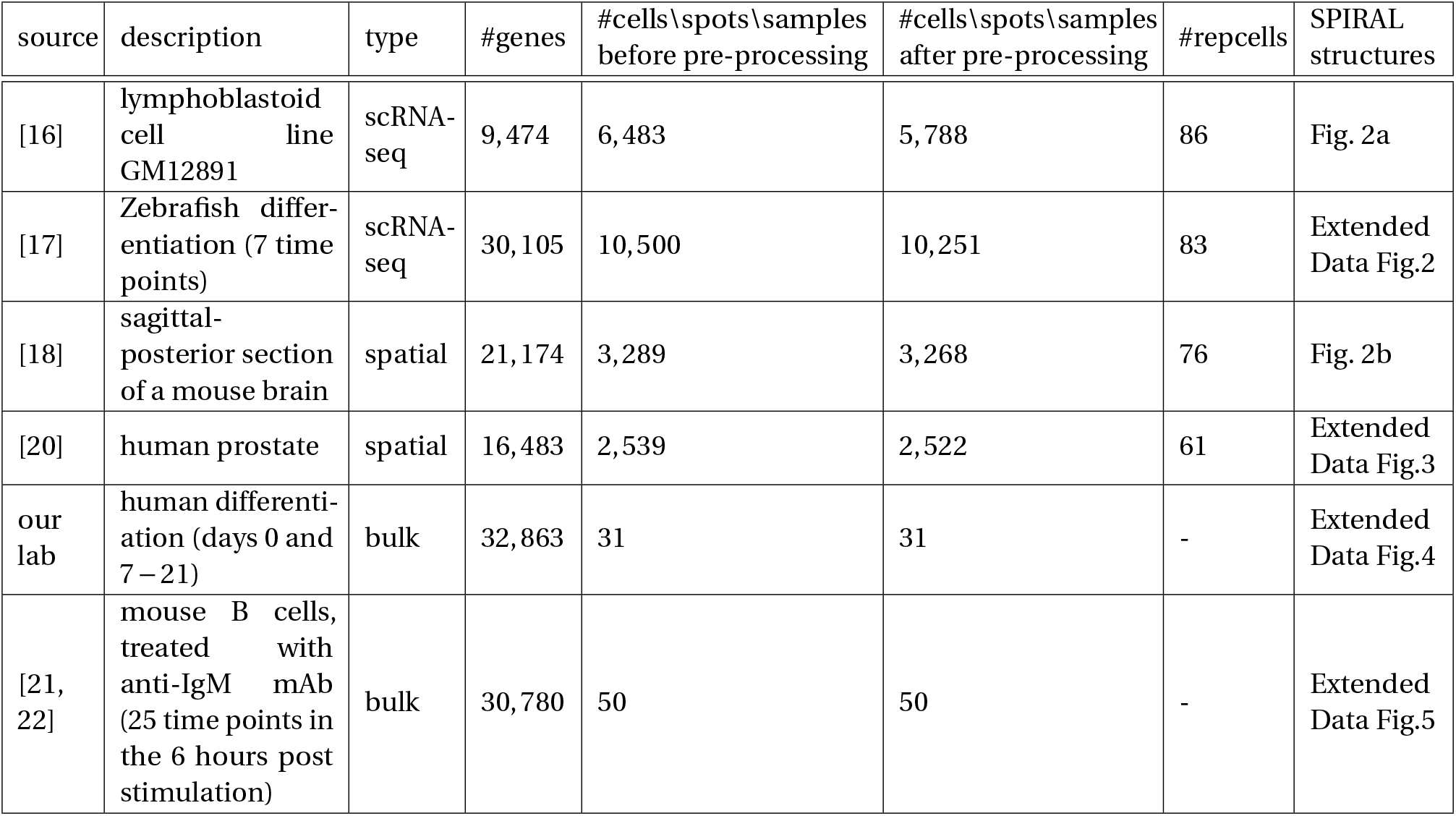
Table 1: datasets

### 9.2 Extended data figure: Zebrafish differentiation single cell data

**Extended Data Fig.2:**
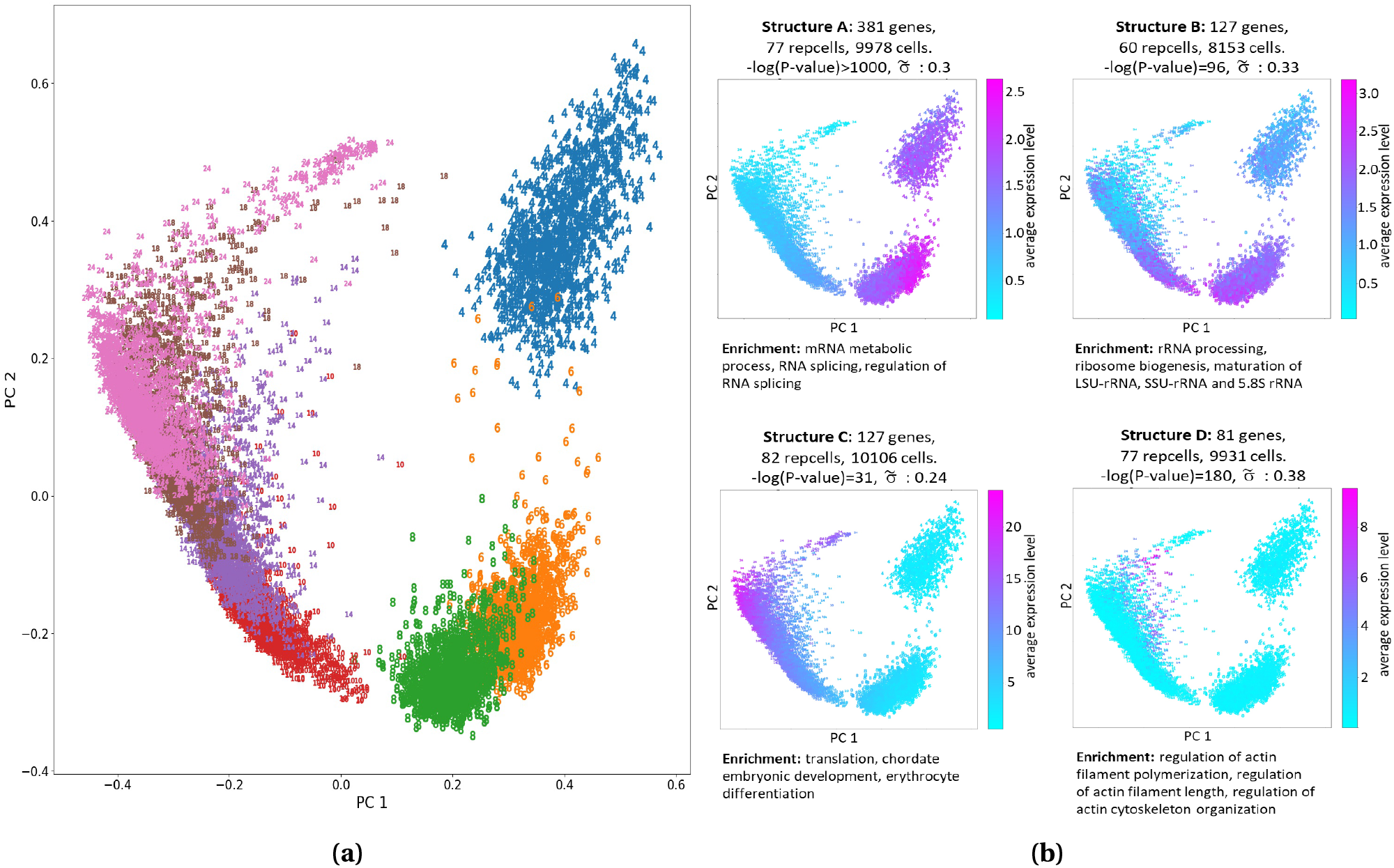
A demonstration of SPIRAL output for a single cell dataset of Zebrafish differentiation [17]. **(a)** PCA layout of raw data. Markers correspond to hours post fertilization. **(b)** PCA layouts of four SPIRAL structures.

### 9.3 Extended data figure: spatial transcriptomics data of a human prostate

**Extended Data Fig.3:**
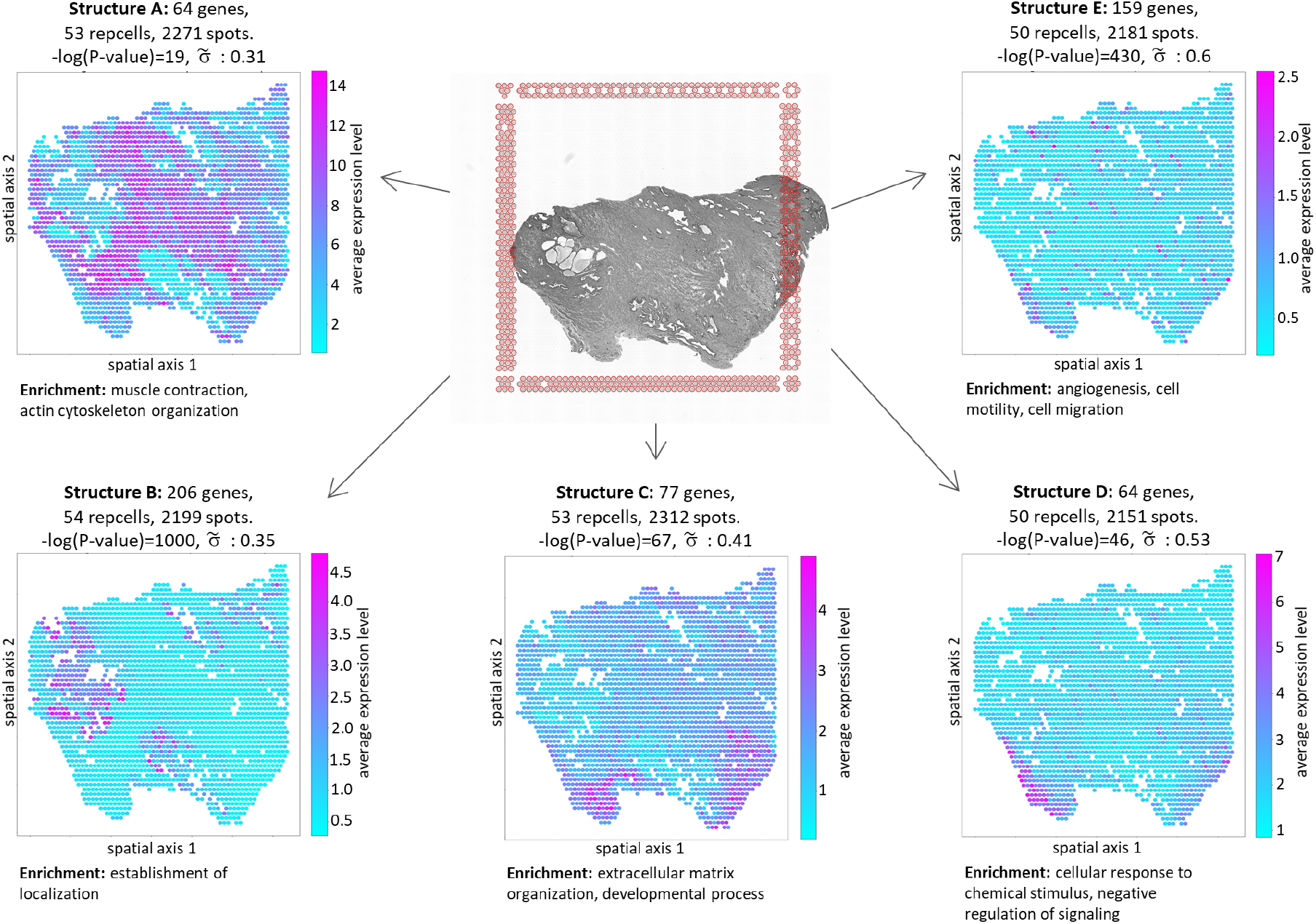
A demonstration of SPIRAL output for a spatial transcriptomics dataset of a normal human prostate [20].

### 9.4 Extended data figure: bulk human differentiation expression data

**Extended Data Fig.4:**
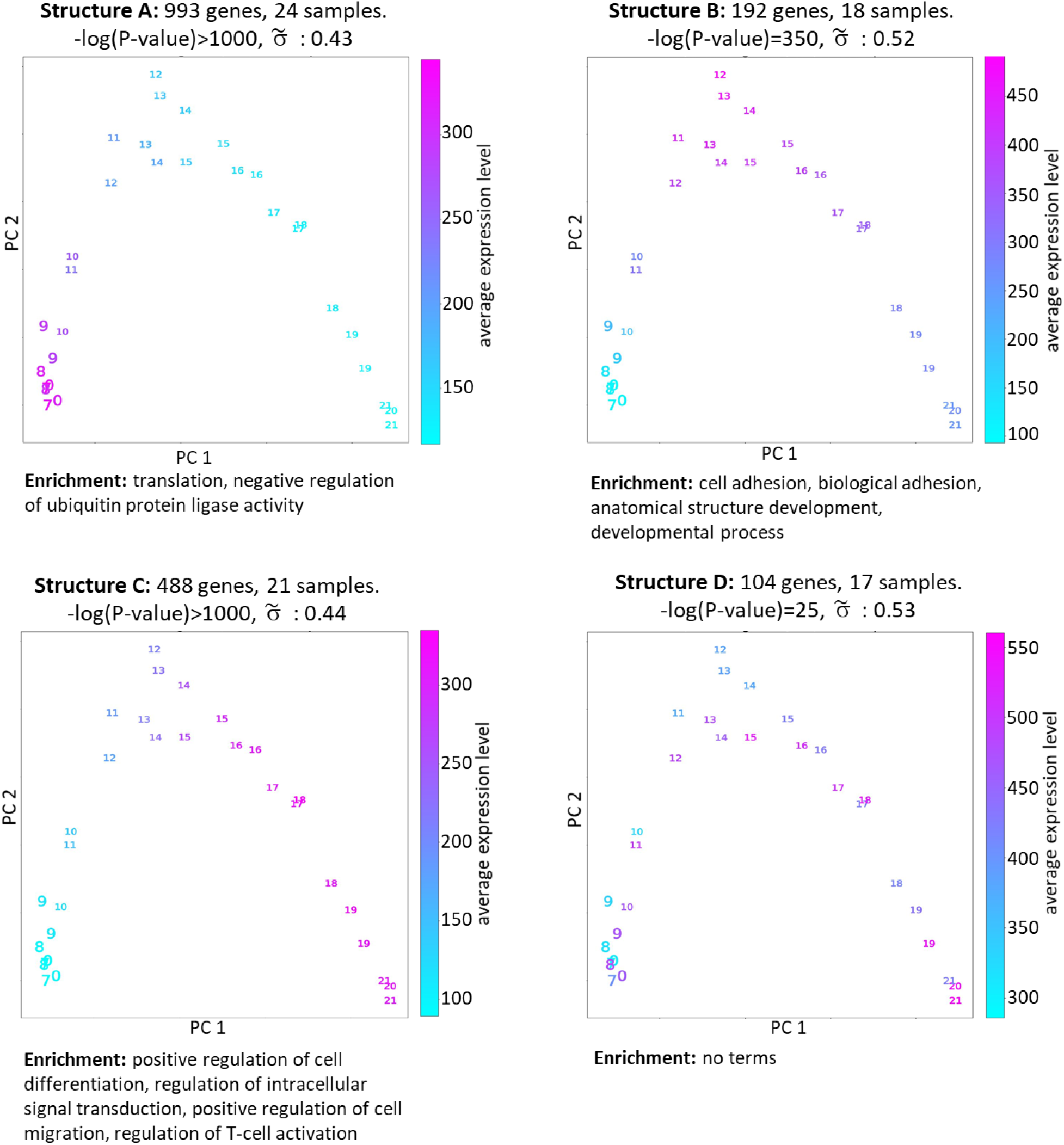
A demonstration of SPIRAL output for a bulk RNA-seq dataset of human embryonic stem cells during differentiation. Markers indicate the day of differentiation.

### 9.5 Extended data figure: bulk expression data of anti-IgM-stimulated mouse B cells

**Extended Data Fig.5:**
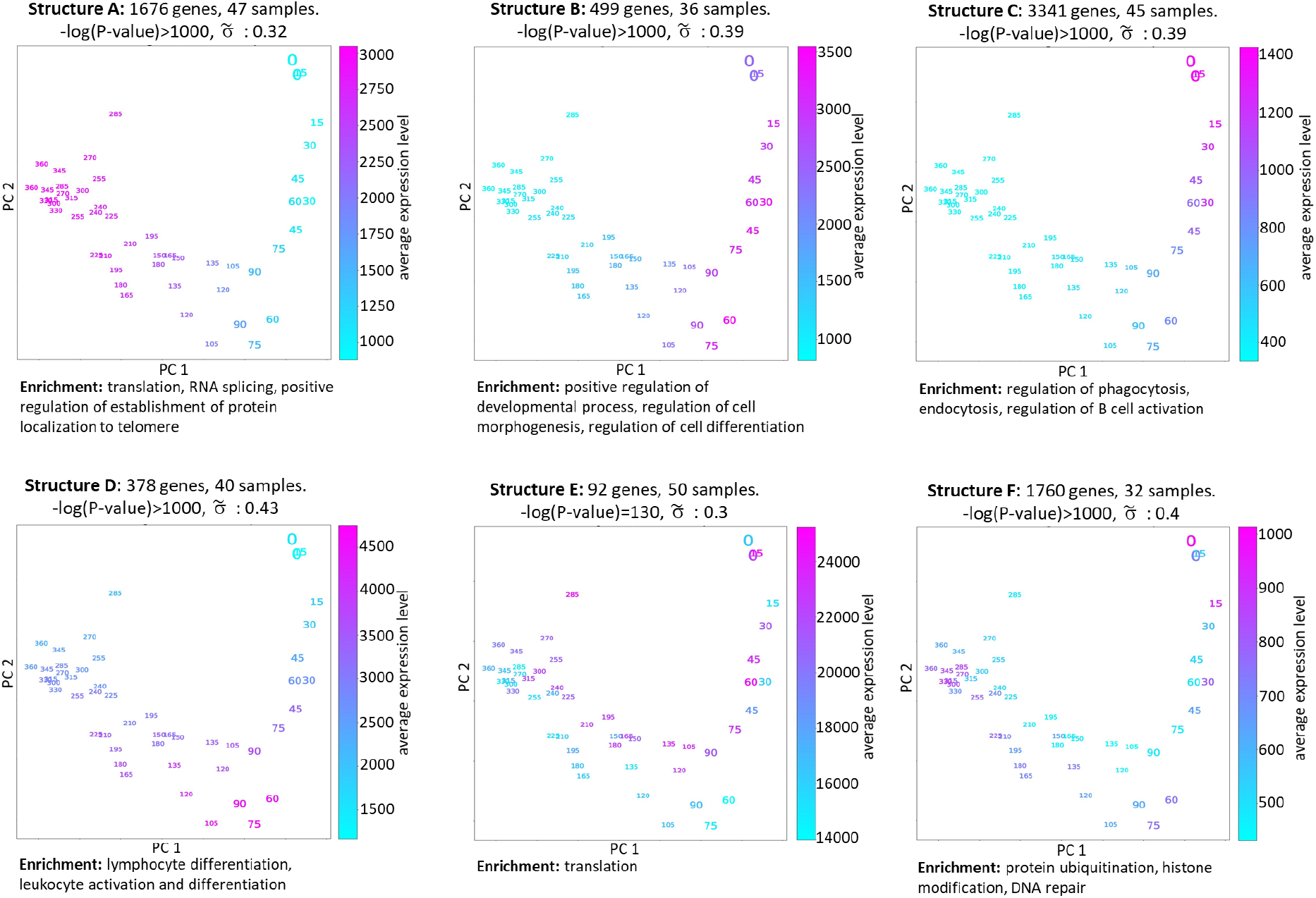
A demonstration of SPIRAL output for a bulk RNA-seq dataset of mouse B cells that were treated with anti-IgM mAb to mimic B cell receptor stimulation. Markers indicate the number of minutes post stimulation.

## Notes

https://spiral.technion.ac.il/

### Competing Interest Statement

The authors have declared no competing interest.

